# Single Cortical Neurons as Deep Artificial Neural Networks

**DOI:** 10.1101/613141

**Authors:** David Beniaguev, Idan Segev, Michael London

## Abstract

We introduce a novel approach to study neurons as sophisticated I/O information processing units by utilizing recent advances in the field of machine learning. We trained deep neural networks (DNNs) to mimic the I/O behavior of a detailed nonlinear model of a layer 5 cortical pyramidal cell, receiving rich spatio-temporal patterns of input synapse activations. A Temporally Convolutional DNN (TCN) with seven layers was required to accurately, and very efficiently, capture the I/O of this neuron at the millisecond resolution. This complexity primarily arises from local NMDA-based nonlinear dendritic conductances. The weight matrices of the DNN provide new insights into the I/O function of cortical pyramidal neurons, and the approach presented can provide a systematic characterization of the functional complexity of different neuron types. Our results demonstrate that cortical neurons can be conceptualized as multi-layered “deep” processing units, implying that the cortical networks they form have a non-classical architecture and are potentially more computationally powerful than previously assumed.

## Introduction

Neurons are the computational building blocks of the brain. Understanding their input-output (I/O) transformation has therefore been a major quest in neuroscience since Ramon y Cajal’s “neuron doctrine”. With the recent development of sophisticated genetical, optical and electrical techniques it has become clear that many key neuron types (e.g., cortical and hippocampal pyramidal neurons, cerebellar Purkinje cells) are highly complicated I/O information processing devices. They receive a barrage of thousands of synaptic inputs via their elaborated dendritic branches; these inputs interact with a plethora of local nonlinear regenerative processes, including the back-propagating (*Na*^+^-dependent) action potential (G. J. Stuart and Sakmann 1994), the multiple local dendritic NMDA-dependent spikes (Polsky, Mel, and Schiller 2004; Branco, Clark, and Häusser 2010; Kastellakis et al. 2015), and the large and prolonged *Ca*^*2*+^ spike at the apical dendrite of L5 cortical pyramidal neurons (M E Larkum, Zhu, and Sakmann 1999). As a result of local nonlinear dendritic processing, a train of output spikes are generated in the neuron axon, carrying information that is communicated, via synapses, to thousands of other (postsynaptic) neurons. Indeed, as a consequence of their inherent nonlinear mechanisms, neurons can implement highly complicated I/O functions (Poirazi, Brannon, and Mel 2003b; Shepherd et al. 1985; C Koch, Poggio, and Torres 1982; Bar-Ilan, Gidon, and Segev 2012; Christof Koch and Segev 2014; London and Häusser 2005; Behabadi and Mel 2013; Häusser and Mel 2003; Mel 1992; Poirazi, Brannon, and Mel 2003a; Moldwin and Segev 2018; Hawkins and Ahmad 2016; Zador, Claiborne, and Brown, n.d.; Bai, Zico Kolter, and Koltun 2019; Cazé, Humphries, and Gutkin 2013; Doron et al. 2017; Greg Stuart, Spruston, and Häusser 2007), and see recent work on dendritic computations in human cortical neurons in (Gidon et al. 2020).

A classical approach to study the I/O relationship of neurons is to ignore their biological details and obtain a highly reduced phenomenological abstraction of the neuron’s I/O characteristics (McCulloch and Pitts 1943; Lapicque 1907). One such abstraction was provided by the “perceptron” (Rosenblatt and F. 1958), which lies at the heart of some of the most advanced pattern recognition techniques to date (LeCun, Bengio, and Hinton 2015) . However, the basic function of the perceptron, a linear summation of its inputs and thresholding for output generation, highly oversimplifies the synaptic integration processes taking place in real neurons. Some studies have addressed this gap, but not at the scale of a realistic number of synaptic inputs, full dendritic biophysical mechanisms, and high temporal precision of spike output (Poirazi, Brannon, and Mel 2003b; Polsky, Mel, and Schiller 2004; Ujfalussy et al. 2018).

Another common approach to study the I/O of neurons is to simulate, via a set of partial differential questions, the fine electrical and anatomical details of the neurons using the cable and compartmental modeling methods introduced by Rall (Rall 1959; Rall 1964; Segev and Rall 1988). Using these models, it is possible to account for nearly all of the above experimental phenomena and to explore conditions that are not accessible with current experimental techniques. However, this success comes at a price because compartmental and cable models are composed of a high dimensional system of coupled nonlinear differential equations, which is notoriously challenging to understand (Strogatz 2001). Specifically, it is a daunting task to extract general principles that govern the transformation of thousands of synaptic inputs to a train of spike output at the millisecond precision from such detailed simulations, but see (Rapp, Yarom, and Segev 1992; G. J. Stuart and Sakmann 1994; G Stuart et al. 1997; Matthew E Larkum et al. 2009; Schiller et al. 2000; Magee and Johnston 1995; Spruston et al. 1995).

Here we propose a novel approach to study the neuron as a sophisticated I/O information processing unit by utilizing recent advances in the field of machine learning. Specifically, we utilized the capability of deep neural networks (DNNs) to learn very complex I/O mappings, in particular that of neurons. Towards this end, we trained DNNs with rich spatial and temporal patterns to mimic the I/O behavior of a layer 5 cortical pyramidal neuron with its full complexity, including its elaborated dendritic morphology, the highly nonlinear local dendritic membrane properties, and a large number of excitatory and inhibitory inputs that bombard the neuron. Consequently, we obtained a computationally efficient DDN model that faithfully predicted the output of this neuron at a millisecond temporal resolution. We then analyzed the weight matrices of the DNN to gain new insights into the I/O function of cortical neurons. By systematically varying the DNN size, this approach allowed us to characterize the functional complexity of a single biological neuron and to pin down its origin. We demonstrated that cortical pyramidal neurons, and the networks that they form, are potentially computationally much more powerful, and “deeper”, than previously assumed.

## Results

Our goal is to fit the I/O relationship of a detailed biophysical neuron model to an analogous DNN. This DNN receives, as a training set, the identical synaptic input and the respective axonal output of the biophysical model. By changing the connection strengths of the DNN using a back-propagation learning algorithm, the DNN should replicate the I/O transformation of the detailed model. To accommodate for the temporal aspect of the input, we employed temporal convolutional networks (TCN) throughout the study.

Figure 1 demonstrates the feasibility and usefulness of this paradigm starting with the I/O transformation of a well understood neuron model: the integrate and fire (I&F) neuron (Lapicque 1907; Burkitt and N. 2006). This neuron receives a train of random synaptic inputs and produces a subthreshold voltage response as well as a spiking output (see **Methods**). But what is the simplest DNN that faithfully captures the I/O properties of this most basic single neuron model? To answer this question, we constructed DNN consisting of one hidden layer with a single hidden unit (Fig. 1**A**). The time axis was divided into 1ms time bins in which only a single spike can occur in the I&F neuron model. The objective of the DNN network is to predict the binary spike output of the I&F model at time t_0_, based on the preceding input spike trains up to t_0_. This input is represented using a binary matrix of size *N*_*syn*_ × *T*, where N_syn_ is the number of input synapses, and *T* is the number of preceding time bins considered (Fig. 1**B**). We used *N*_*syn*_ = 100, and trained a DNN with a single hidden unit on 5000 seconds of simulated data. When using T = 80ms we achieved a very good fit, namely: a simple DNN with a single hidden unit that accurately predicted both the subthreshold voltage dynamics as well as the spike output of the respective I&F neuron model at a millisecond precision (Fig. 1**C**).

**Fig. 1.**
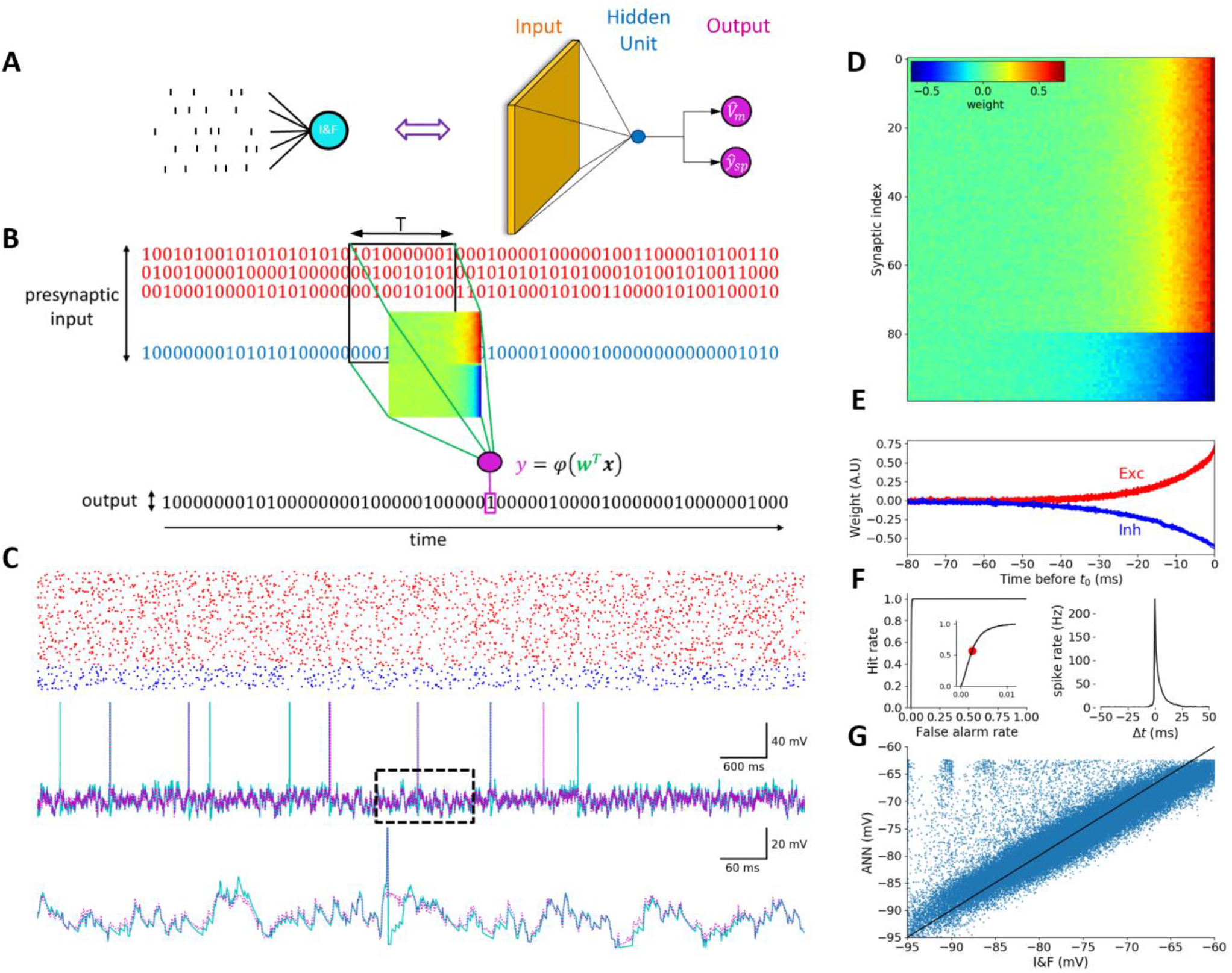
Integrate and Fire neuron model is faithfully captured by a DNN with one hidden layer consisting of a single hidden unit. (**A**) Illustration of an I&F neuron model receiving a barrage of synaptic inputs and generating voltage/spiking output (left), and its analogous DNN (right). Orange, blue, and magenta represent the input layer, the hidden layer and the DNN output, respectively. (**B**) Schematic overview of our prediction approach. The objective of the DNN is to predict the spike output of the respective I&F model based on its synaptic input. The binary matrix, denoted by **x**, represents the input spikes in a time window T (black rectangle) preceding t_0_. **x** is multiplied by the synaptic weight matrix, **w** (represented by the heatmap image) and summed up to produce the activation value of the output unit **y**. This value is used to predict the output (magenta rectangle) at t = t_0_. Excitatory input is denoted in red; inhibitory in blue. Note that, unlike the I&F, the DNN has no *a priori* information about the type of the synaptic inputs (E or I). (**C**) **Top**. Example inputs (red – excitatory, blue – inhibitory) presented to the I&F neuron model. **Middle**. Response of the I&F model (cyan) and of the analogous DNN (magenta). **Bottom**. Zoom in on the dashed rectangle region in the top trace. Note the great similarity between the two traces. (**D**) Learned weights of the DNN modeled synapses. Top 80 rows are excitatory synapses to the I&F model; bottom 20 rows are its inhibitory synapses. Columns correspond to different time points relative to t_0_ (right most time point). The prediction probability for having a spike at t_0_ increases if the number of active excitatory synapses increases (red) and the number of active inhibitory synapses decreases (blue) just before t_0_. (**E**) Temporal cross section of the learned weights in **D**. (**F**) **Left**. Receiver Operator Characteristic (ROC) curve of spike prediction. The area under the curve (AUC) is 0.997, indicating high prediction accuracy at 1ms precision. Inset: zoom in on up to 1% false alarm rate. Red circle denotes the threshold selected for the DNN model shown in **C**. Right. Cross Correlation between the I&F spike train (ground truth) and the predicted spike train of the respective DNN, when the prediction threshold was set to 0.2% false positive (FP) rate (red circle in left plot). (**G**) Scatter plot of the predicted DNN subthreshold voltage versus ground truth voltage produced by the I&F model).

Figure 1**D** depicts the weights (“filters”) of the single hidden unit of the respective DNN as a heatmap. It shows that the learning process automatically produced two classes of weights (“filters”), one positive and one negative, corresponding to the excitatory and inhibitory inputs impinging on the I&F model. In agreement with our understanding of the I&F model, the excitatory inputs contribute positively to output spike prediction (red color), whereas the inhibitory inputs contribute negatively it (blue color). Earlier inputs, either inhibitory or excitatory, contribute less to this prediction (teal color). Figure 1**E** depicts the temporal cross-section of those filters and reveals an exponential profile that reflects the temporal decay of post synaptic potentials in the I&F model (in the reverse time direction). From these filters one can recover the precise membrane time constant of the I&F model. These two temporal filters (excitatory and inhibitory) are easily interpretable as they agree with our previous understanding of the temporal behavior of synaptic inputs that give rise to an output spike in the I&F model.

Figure 1**F** quantifies the model performance in terms of spike prediction using the Receiver Operating Characteristic (ROC) curve (Fig. 1**F** left and see **Methods**) and the area under it (AUC). AUC for the I&F case is 0.9971, indicating a very good fit. Figure 1**F** right shows an additional quantification of spike temporal precision using the DNN prediction, by plotting the cross correlogram between the predicted spike train and the target I&F simulated spike train (the “ground truth”). The cross correlogram shows a sharp peak at 0 millisecond and has a short, ~10 millisecond, half-width, suggesting high temporal accuracy of the DNN. We also quantified the DNN performance to predict the subthreshold membrane potential by using standard regression metrics, and, in Fig 1**G**, depict the scatter plot of the predicted voltage versus the ground truth simulated output voltage. The Root Mean Square Error (RMSE) is 1.23mV, indicating a good fit between the I&F and the respective DNN.

In conclusion, as a proof of concept, we have demonstrated that a simple DNN can learn the I/O transformation of a simple I&F model with a high degree of temporal accuracy. The weight matrix (the “filter”) obtained by the learning process represents known features of the I&F model, including the existence of two classes of inputs (excitatory and inhibitory), the convolution of the synaptic inputs with the exponential decay representing the passive membrane RC properties, and the transformation from subthreshold membrane potential to spike output.

We next applied our paradigm to a morphologically and electrically complex detailed biophysical compartmental model of a 3D reconstructed layer 5 cortical pyramidal cell (L5PC) from rat somatosensory cortex (Fig. 2**A**). The model is equipped with complex nonlinear membrane properties, a somatic spike generation mechanism and an excitable apical nexus capable of generating calcium spikes (Hay et al. 2011; M E Larkum, Zhu, and Sakmann 1999). The excitatory synaptic inputs are mediated through both voltage-independent AMPA-based conductance as well as by voltage-dependent NMDA-type conductance; the inhibitory inputs are mediated through conductance-based GABA_A_-type synapses. Both excitatory and inhibitory synapses are uniformly distributed across the dendritic tree of the model neuron (see **Methods**).

**Fig 2.**
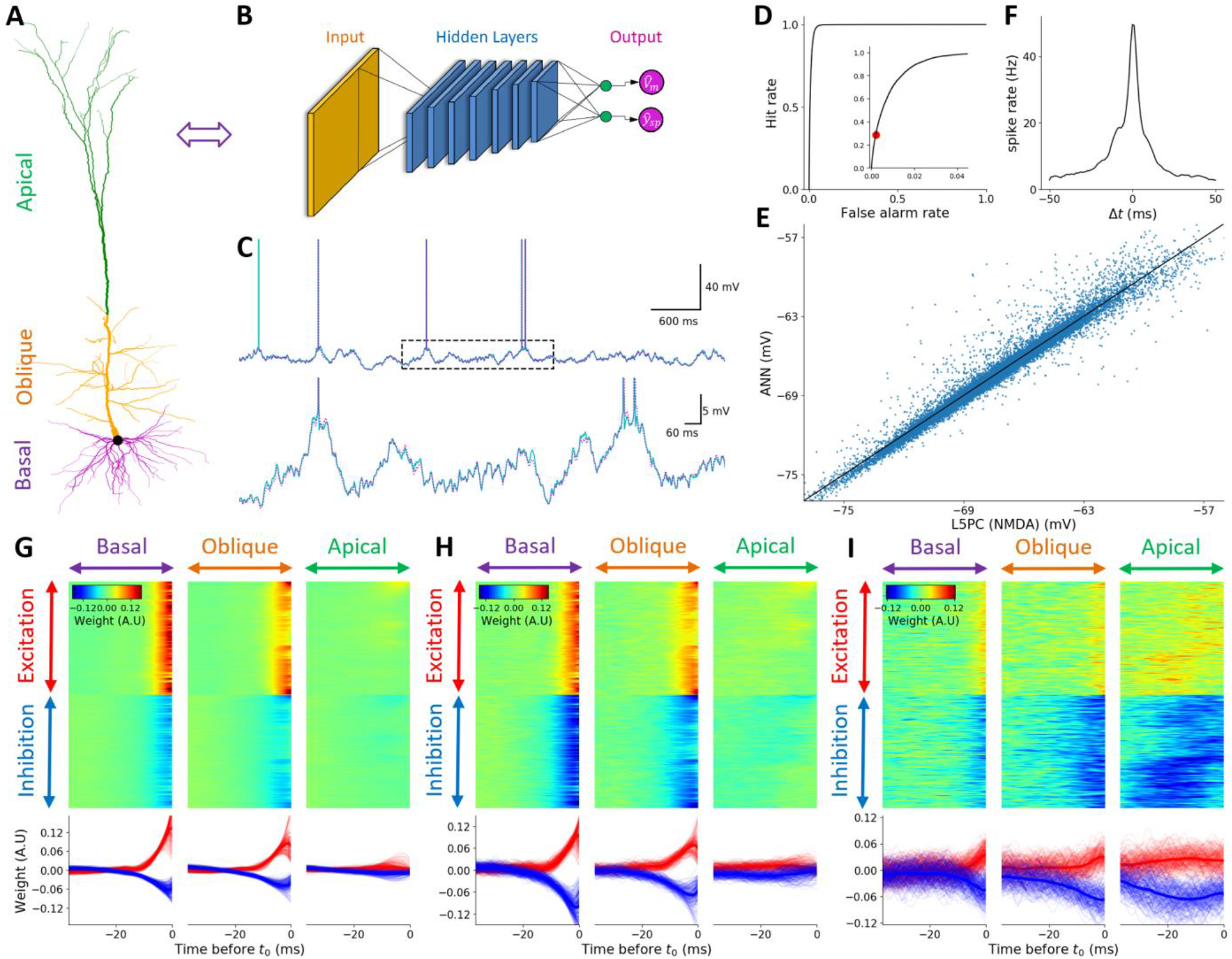
Detailed model of a layer 5 pyramidal neuron is faithfully captured by a TCN with 7 hidden layers consisting of 128 feature maps at each layer. (**A**) The modeled L5PC. Basal, oblique, and apical dendrites are marked purple, orange, and green, respectively. (**B**) analogous DNN. Orange, blue, and magenta circles represent the input layer, the hidden layer, and the DNN output, respectively. Green units represent linear activation units (see Methods). (**C**) Top. Exemplar voltage response of the L5PC model with NMDA synapses (cyan) and of the analogous DNN (magenta) to random synaptic input stimulation. Bottom. Zoom in on the dashed rectangle region in the top trace. (**D**) ROC curve of spike prediction; the area under the curve (AUC) is 0.9911, indicating high prediction accuracy at 1ms precision. Zoom in on up to 4% false alarm rates is shown in the inset. Red circle denotes the threshold selected for the DNN model shown in **B**. (**E**) Scatter plot of the predicted DNN subthreshold voltage versus ground truth voltage. (**F**). Cross Correlation plot between the ground truth (L5PC with NMDA synapses) spike train and the predicted spike train of the respective DNN, when the prediction threshold was set according to red circle in **D**. (**G**) Learned weights of a selected unit in the first layer of the DNN. Top Left, top center and top right, inputs located on the basal dendrites, on the oblique dendrites and on the apical tuft, respectively. For each case, top half of the rows are excitatory synapses whereas bottom half of rows are its inhibitory synapses. Different columns correspond to different time points relative to t_0_ (rightmost time point). Bottom. temporal cross-section of the learned weights above. (**H**) Similar to **G**, first layer weights for a different unit in the first layer but with a different spatio-temporal pattern. (**I**) An additional unit that is weakly selective to whatever happens in the basal dendrites, weakly sensitive to oblique dendrites, but very sensitive to apical tuft dendrites. The output of this hidden unit is increased when there is apical excitation and lack of apical inhibition in a time window of 40 ms before t_0_. Note the asymmetry between the amplitudes of the temporal profiles of excitatory and inhibitory synapses, indicating that inhibition decreases the activity of this unit more than excitation increases it.

A thorough search of configurations of deep and wide fully-connected neural network architectures (FCNs) have failed to provide a good fit to the I/O characteristics of the L5PC model. These failures suggest a substantial increase in the complexity of I/O transformation compared to that of I&F. Indeed, only temporally convolutional network architecture (TCN) with 7 layers and 128 channels per layer, provided a good fit (Fig. 2**B, C** Fig. S5). The example in Fig. 2C shows that this TCN, when provided with a previously unseen input pattern from the test set, can predict the somatic voltage and spikes of a highly complex neuron with high precision.

It is important to note that the accuracy of the model was insensitive to the temporal kernel sizes of the different DNN layers when keeping the total temporal extent of the entire network fixed, so the temporal extent of the first layer was selected to be larger than subsequent layers mainly for visualization purposes (see Fig. 2**G**-**I**). Figure 2**H** shows a filter from a unit at the first layer of the DNN. This filter is somewhat similar to the filter in Fig. 1**D** but integrates only basal and oblique subtrees and ignores the inputs from the apical tree. Figure 2**I**, however, shows a filter of another unit that, in contrast to Fig. 2**H**, has negligible weights assigned for basal and oblique dendrites but a very strong apical tuft dependency. By examining additional first layer filters (not shown) we found a wide variety of different activation patterns that the TCN utilized as an intermediate representation, including many temporally directionally selective filters (similar to those of Fig. 4**D**). Figure 2**D**,2**E**,2**F** shows the quantitative performance evaluation of this DNN model. For binary spike prediction (Fig. 2**D**), the AUC is 0.9911. For somatic voltage prediction (Fig. 2**E**), the RMSE is 0.71mV and 94.6% of the variance is explained by this model. Note that, despite its seemingly large size, the resulting TCN represents a substantial decrease in computational resources relative to a full simulation of a detailed biophysical model (involving numerical integration of thousands of non-liner differential equations), as indicated by a speedup of simulation time by several orders of magnitude.

Now that we have obtain an alternative DNN model that can replicate very accurately the I/O relationship of a detailed biophysical/compartmental model of a real neuron, can we learn from it what are the essential features that contribute to neuron complexity? Detailed studies of synaptic integration in dendrites of cortical pyramidal neurons suggested the primary role of the voltage-dependent current through synaptic NMDA receptors, including at the subthreshold and suprathreshold (the NMDA-spike) regimes (Polsky, Mel, and Schiller 2004; Branco, Clark, and Häusser 2010). As NMDA receptors depend nonlinearly on voltage it is highly sensitive not only to the activity of the synapse in which the receptors are located but also to the activity of (and the voltage generated by) neighboring synapses and to their dendritic location. Moreover, the NMDA-current has slow dynamics, promoting integration over a time window of tens of milliseconds (Major, Larkum, and Schiller 2013; Doron et al. 2017). Consequently, we hypothesized that removing NMDA dependent synaptic currents from our L5PC model will significantly decrease the size of the respective DNN.

Fig. 3 shows that, after removing the NMDA voltage dependent conductance, such that the excitatory input relies only on AMPA mediated conductances, we have managed to achieve a similar quality fit as in Fig. 2 when using a much smaller network - a fully connected DNN (FCN) with 128 hidden units and only a single hidden layer (Fig. 3**B**). This significant reduction in complexity is due to the ablation of NMDA channels. Also, in our DNN training attempts, we have failed to achieve a good fit when using the architecture that was successful for the I&F model neuron shown in Fig. 1. This indicates that, whereas the DNN model for L5PC is greatly simplified in the absence of NMDA conductance, additional neuronal mechanisms still contribute to the richness of its I/O transformation as compared to that of the I&F neuron model.

**Fig 3.**
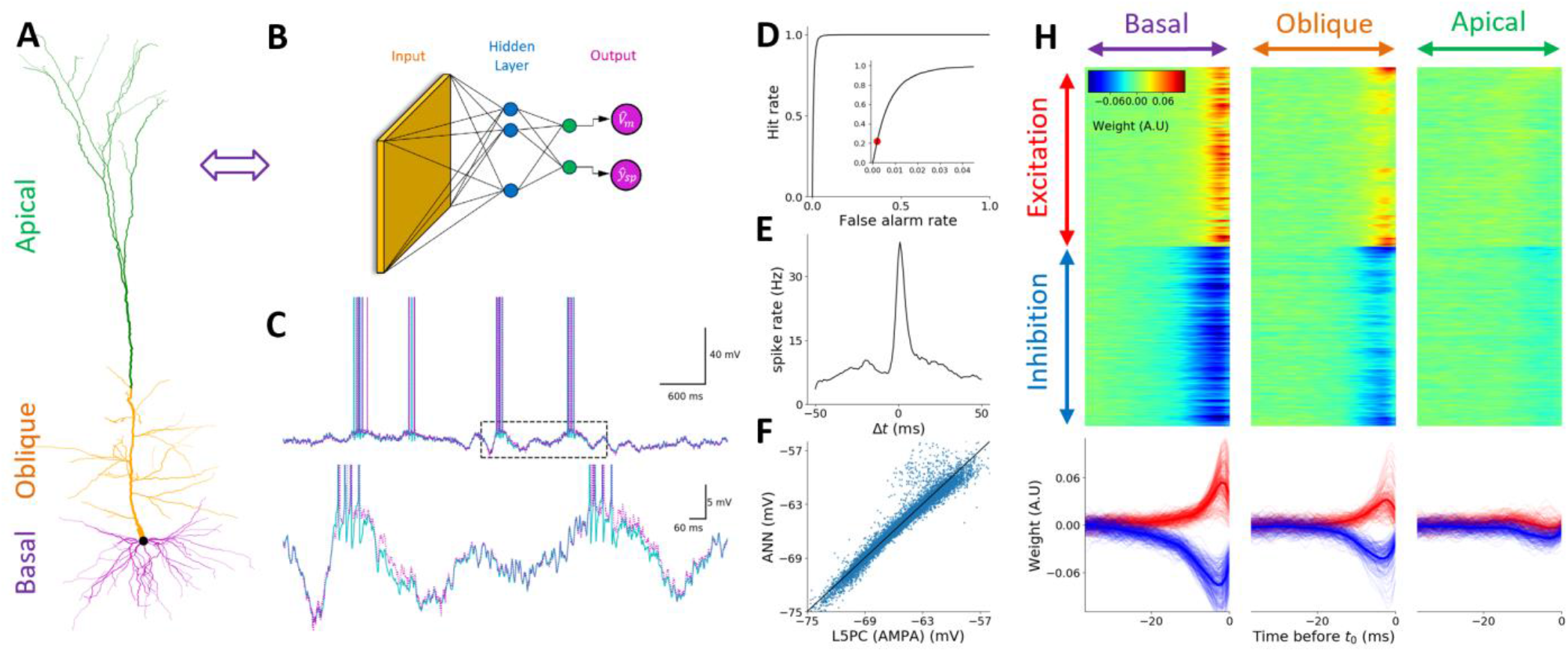
Detailed model of L5PC neuron without NMDA synapses is faithfully captured by DNN with one hidden layer consisting of 128 hidden units. (**A**) Illustration of L5PC model (**B**) analogous DNN. As in previous figure, orange, blue and magenta circles represent the input layer, the hidden layer and the DNN output, respectively. Green units represent linear activation units. (**C**) Top. Response of the L5PC model (cyan) and of the analogous DNN (magenta) to random AMPA-based excitatory and GABAA-based inhibitory synaptic input (see **Methods**). Bottom. Zoom in on the dashed rectangle region in the top trace. Note the great similarity between the two traces. (**D**). ROC curve of spike prediction; the area under the curve (AUC) is 0.9913, indicating high prediction accuracy at 1ms precision. Inset: zoom in on up to 4% false alarm rates. Red circle denotes the threshold selected for the DNN model shown in **B**. (**E**) Cross Correlation plot between the ground truth (L5PC model response) and the predicted spike train of the respective DNN, for prediction threshold indicated by red circle in left plot. (**F**) Scatter plot of the predicted DNN subthreshold voltage versus ground truth voltage. (**H**) Learned weights of selected units in the DNN, separated by their morphological (basal, oblique and apical) location. Like in the previous figure, in each case, the top half rows are excitatory synapses and the bottom half are the inhibitory synapses. As in Fig 1**D**, different columns correspond to different time points relative to t_0_ (rightmost time point). Note that, just before t_0_, the output of this hidden unit increases if the number of active excitatory synapses increases at the basal and oblique dendrites (red), whereas the number of active inhibitory synapses decreases (blue) at these locations. However, this unit is non-selective to activity at the apical tuft, indicating the lack of influence of the tuft synapses on the neuron’s output.

Fig. 3**C** shows an exemplar test trace for the DNN illustrated in Fig. 3**B**, whereas **Fig.** 3**H** depicts a representative exemplar of the weight matrix for one of the hidden units of the DNN. By examining the filters of the hidden layer of the DNN we observed that the weights representing inputs to the oblique and basal dendrites had profiles that resembles PSPs (but mirrored in time). Interestingly, the weights to the apical tuft are essentially zero. This pattern remains consistent for all first layer filters of the network, implying that, for this model, the apical dendritic synapses had negligible information regarding predictions of the output spikes of the neuron, even in the presence of calcium spikes occasionally occurring in the nexus. Contrasting this filter with the one presented in Fig. 2**I**, suggests that the NMDA non-linearity greatly assists in the activation of apical tuft dendrites.

Fig. 3**D**, 3**E** and 3**F** show the quantitative performance evaluation. For binary spike prediction (Fig. 3**E**), the AUC is 0.9913. For somatic voltage prediction (Fig. 3**F**), the RMSE is 0.58mV and 95.0% of the variance is explained by this model.

To further investigate the NMDA contribution to computational complexity, we studied it in isolation on a single dendritic branch and attempted to fit the somatic voltage of a Layer 2/3 Pyramidal Cell (L23PC), taken from the visual cortex of the mouse (Branco, Clark, and Häusser 2010), in response to random activation of only 9 excitatory synapses uniformly distributed across a single dendritic branch (Fig. 4**A**). We found that a single dendritic branch with NMDA synapses is faithfully captured by a single layer of a fully connected DNN with 4 hidden units (Fig. 4**A**&**B**). Examining the 4 filters of the first layer reveals interesting shapes that make intuitive sense (as first explored by the pioneering theoretical studies of (Goldstein and Rall 1974; Rall 1969; Rall 1964)). The topmost filter in Fig. 4**D** appears to be summing only very recent and proximal dendritic activation. The second-from-top hidden unit sums up recent distal dendritic synaptic inputs. The third filter clearly shows a direction selective hidden unit, preferring patterns in which synaptic activation are temporally activated from distal to proximal series, and the last hidden unit responds to a prolonged distal dendrite summation of activity combined with precisely timed proximal input activation.

**Fig 4.**
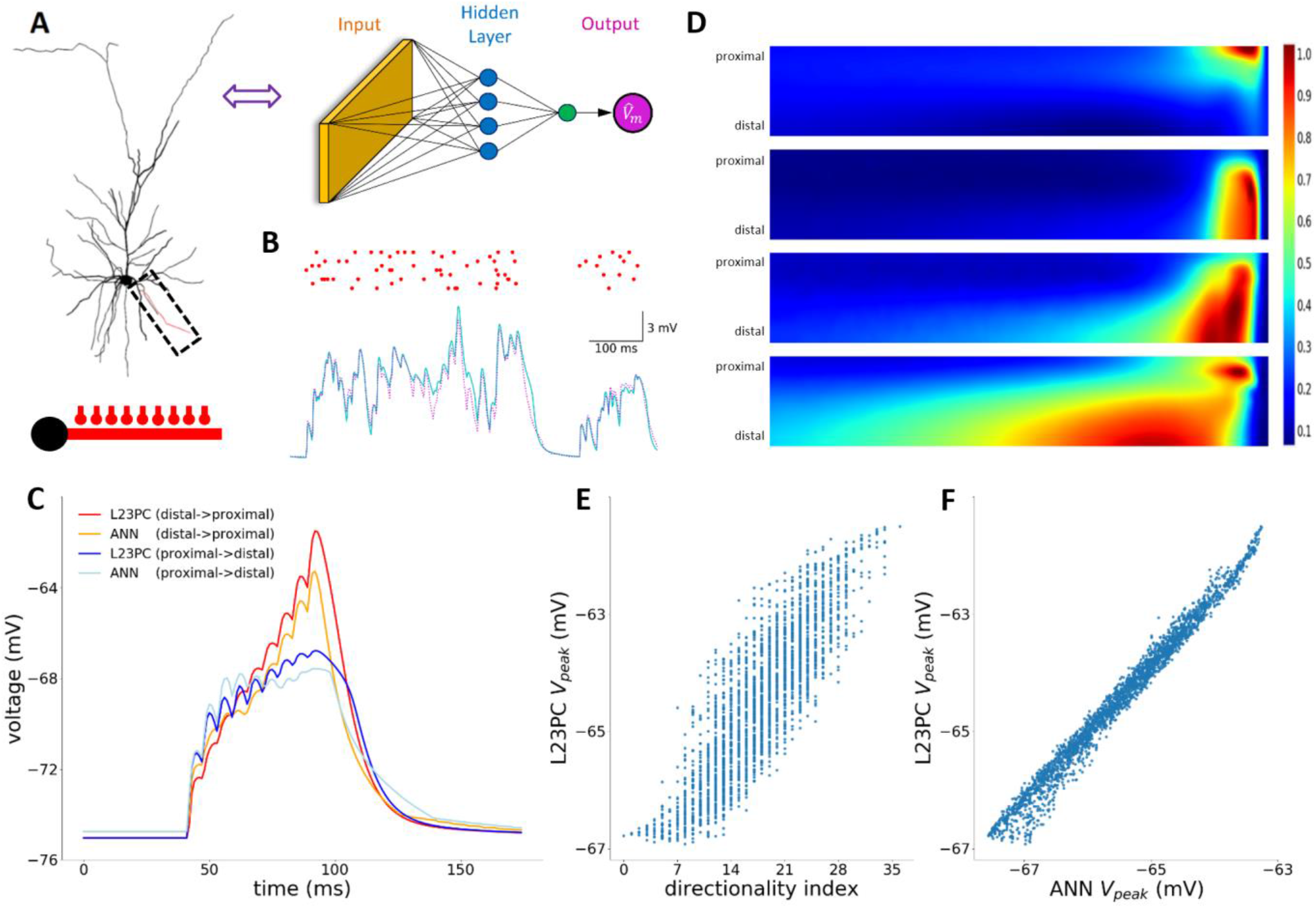
Response of a single dendritic branch of L2/3PC neuron model receiving spatio-temporal pattern of NMDA synapses activation is faithfully captured by a DNN with one hidden layer consisting of 4 hidden units. (**A**) Left. Layer 2/3 pyramidal neuron used in the simulations with a zoom in on one selected basal branch (dashed rectangle). Same modeled branch with 9 excitatory synapses (depicted schematically by the “ball and stick” at bottom) was also used in the study of Branco et al. (Branco, Clark, and Häusser 2010). Right. Illustration of the analogous DNN. Colors as in Fig. 2**A**. (**B**) Example of the somatic voltage response (cyan) and DNN predicted output (magenta) to a randomly generated input spike pattern on that basal branch (red dots above). (**C**) Example of somatic response to two spatio-temporal sequences of synaptic activation patterns (red: distal-to-proximal direction” and blue: “proximal-to-distal direction”) and the DNN predicted output for these same sequences (orange and light blue traces, respectively). (**D**) Learned weights of the 4 hidden units by the respective DNN model. Heatmaps are spatio-temporal filters as shown in Figs. 1**D** and 2**C**. Note the direction selective shapes and long temporal extent of influence by distal synaptic activations. (**E**) Scatter plot that shows the discrimination ability between different orders of synaptic activations on the modeled basal branch. Vertical axis is the ground truth maximum voltage at the soma during a specific synaptic order of activation. Horizontal axis is directionality index proposed in Branco et al. (Branco, Clark, and Häusser 2010). Correlation coefficient is 0.86. (**F**) Same as **E**, but the DNN estimation of the max voltage of the respective order of activation, showing a superior performance of the DNN prediction relative to the directionality index previously proposed. Correlation coefficient is 0.99.

Fig. 4**C** specifically examines the special cases studied in (Branco, Clark, and Häusser 2010), of sequential activation of the 9 synapses in the distal to proximal direction and, conversely, in the proximal to distal direction. Our DNN network, trained on random synaptic activation patterns, successfully replicated this behavior. Fig. 4**E** shows our reconstruction of the results of Branco et al. (Branco, Clark, and Häusser 2010), whereby a directionality index was suggested as a possible predictor for the peak somatic voltage for random activation sequence of the 9 input synapses. Fig. 4**F** shows the prediction of the respective DNN for the same sequences as in Fig. 4**E**. Interestingly, our DNN acts as a much better predictor for the peak somatic voltage. It is important to note that the special case of 9 synaptic activations equally spaced in time is highly unlikely to occur during the random input stimulation regime which was used to train the DNN. Nevertheless, the network can generalize even to this new input regime with high precision.

These results demonstrate the power of our approach in interpretability and ability of our models to generalize to previously unseen input patterns (out of distribution generalization). By examining the four kernels we provide an intuitive (Fig. 4D), yet powerful (Fig. 4F), interpretation for the complex process of nonlinear synaptic integration in a single dendrite with NMDA synapses. In addition, the network generalizes and predicts the outputs for temporal patterns very different from those on which it was trained.

## Discussion

Recent advances in the field of deep neural networks (DNNs), provide, for the first time, a powerful general-purpose tool that can learn a mapping of I/O relationships from examples, including that of single complex nonlinear neurons, as demonstrated by the present work. We constructed a large dataset of pairs of input-output examples by simulating a neuron model receiving a rich repertoire of synaptic inputs and recorded its output in terms of spikes at millisecond precision as well as its subthreshold membrane potential. We then trained networks of various configurations on these input-output pairs until we obtained an analogous “deep” network with close performance to that of the neuron’s detailed simulation. We applied this framework to a series of neuron models with various levels of morpho-electrical complexity.

For simple I&F neuron models, our framework provides simple DNNs that capture the full I/O relationship of the model while providing key insights that are consistent with our understanding of the I&F models (Fig. 1). Surprisingly, even a neuron model of a layer 5 cortical pyramidal neuron with complex dendritic trees and with a host of dendritic voltage dependent currents and AMPA-based synapses is well-captured by a relatively simple network with a single hidden layer (Fig. 3). However, in a full model of a L5 pyramidal neuron consisting of NMDA-based synapses, the complexity of the network is significantly increased, and we found a good fit with a temporal-convolutional network that has 7 hidden layers (Fig. 2, see also Fig. S4 & S5). However, in the case where there wasonly a single dendritic branch consisting of NMDA synapses, a shallow (one hidden layer) DNN with only a few units was required to capture different aspect of spatio-temporal integration of synaptic inputs (Fig. 4).

These results suggest that the single cortical neuron with its nonlinear synaptic inputs is already, on its own, a sophisticated computational unit. Consequently, cortical networks are “deeper” and computationally more powerful than they seem to be just based on their anatomical (pre-to-post synaptic) connections. Importantly, the implementation of the I/O function using a DNN also provides practical advantages as it is much more efficient than the traditional compartmental model. In our tests we obtained speed up of several orders of magnitude when using the DNN instead of its compartmental-model counterpart. This could make an important contribution to simulating large scale realistic neuronal networks (Markram et al. 2015; Egger et al. 2014). Furthermore, the size of the respective DNN for a given neuron could be used (under certain assumptions, see below) as an index for its computational power; the larger it is the more sophisticated computations this neuron could perform. Such an index will enable a systematic comparison between different neuron types (e.g., CA1 pyramidal cell, cortical pyramidal cell, and Purkinje cell, or for the same type of cell in different species, e.g., mouse vs. human cortical pyramidal cells).

It is important to emphasize that, due to optimization, the complexity measure described above is an upper bound of the true computational complexity of the I/O of a single neuron, i.e., it is possible that there exists a much smaller neural network that could mimic the biophysical neuron with a similar degree of accuracy but the training process we used could not find it. Additionally, we note that we have limited our architecture search space only to fully connected (FCN) and temporally convolutional (TCN) neural network architectures. It is likely that additional architectural search could yield even simpler and more compact models for any desired degree of prediction accuracy. In order to facilitate this search inthe scientific community, we hereby release our large readymade dataset of simulated inputs and outputs of a fully complex single layer 5 cortical neuron in an *invivo* like regime so that the community can focus on modelling various aspects of this endeavour and avoid running the simulations themselves.

The analysis of deep neural networks is a challenging task and a rapid growing field (Olah, Mordvintsev, and Schubert 2017; Mordvintsev, Olah, and Tyka 2015; Mahendran and Vedaldi 2015). Nevertheless, observing the weight matrix of units (“filters”) in first layer of the respective DNN is straightforward and can provide ample insights regarding the I/O transformation of the neuron. The full network can be interpreted as consisting of a basic set of filters that span the space of possible spatio-temporal patterns of synaptic inputs that will drive the original neuron to spike. The first layer defines this space, and the rest of the network mixes and matches within that space. For example, as shown in Fig. 3, in the case of a pyramidal neuron without NMDA synapses, most filters have significant weights only for basal and oblique inputs, and the weight given for apical tuft synapses is negligible (despite the existence of voltage dependent Ca^2+^ and other nonlinear currents in this model including occasional Ca^2+^ spikes). The picture is fundamentally different when NMDA synapses were included in the model. In this case the weights assigned to apical dendrite synapses is significant. Moreover, the filters devoted to these apical inputs tend to have a temporal structure that is significantly wider (in time) than for the proximal synapses, suggesting that the temporal precision of input to the apical synapses is less important. These are basic insights that could be drawn by just observing first layer filters of the resulting DNN.

This work opens multiple additional avenues for future research. One important direction is the isolation of the contribution of specific mechanisms to the computational power of the neuron in a similar way to that performed here for NMDA (See also Fig. S5 & S7). By fitting a DNN while manipulating specific dendritic currents (e.g., VGCa, Kv, HCN channels), we will better understand their contribution to the overall synaptic integration process and to the complexity of the respective DNN. An additional interesting direction is to utilize this work to improve our understanding of dendritic integration. Rather than modelling a neuron bombarded with random meaningless input, one could utilize the neuron’s equivalent-DNN to make the neuron compute some interesting meaningful functions. For example, now that we estimate that a cortical L5 pyramidal neuron is equivalent to a deep network with 7 hidden layers, this DNN could be used to teach the respective neuron to implement a function which is in the scope of the capabilities of such a network, such as classifying hand written digits or a sequence of auditory sounds. One can then both validate the hypothesis that single neurons could perform complex computational tasks and investigate how these neurons can implement such complex tasks.

If one cortical neuron is equivalent to a multi-layered DNN, what are the implications for the cortical microcircuit? Is that circuit merely a “deeper” DNN composed of simple “point neurons”? In fact, a key difference is that, under our model, synaptic plasticity can take place only in the synaptic layer (input) of the respective DNN for a single cortical neuron, whereas the weights of the hidden layers are fixed (and dedicated to represent the I/O function of that single cortical neuron). Taken together with the myriad of recurrent connections and network motifs between cortical neurons of different types (Markram et al. 2015), we hereby propose a very specific network architecture for cortical networks. Indeed, the architecture of artificial neural networks is one of the most rewarding avenues of machine learning today (He et al. 2015; Vaswani et al. 2017; Lin, Chen, and Yan 2014), and studying the specific architecture suggested by our work may unravel some of the inductive bias hidden within the cortical microcircuit and harness it for future AI applications.

## Methods

### I&F simulation

For Fig.1 simulations, membrane voltage was modeled using a leaky I&F simulation 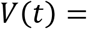 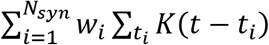, where *w*_*i*_ denotes synaptic efficacy for each synapse, *t*_*i*_ denotes presynaptic spike times, and *K*(*t* − *t*_*i*_) denotes the temporal kernel of each postsynaptic potential (PSP). We used a temporal kernel with exponential decay 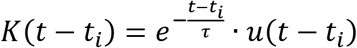 where *u*(*t*) is the Heaviside function 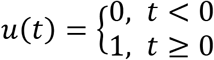 and *τ* = 20*ms* is the membrane time constant. When threshold was reached, an output spike was recorded, and the voltage was reset to *V*_*rest*_ = −95*mV*. As input to the simulated I&F neuron, *N*_*exc*_ = 80 excitatory synapses and *N*_*inh*_ = 20 inhibitory synapses were used. Synaptic efficacies of *w*_*exc*_ = 5*mV* were used for excitatory synapses and *w*_*inh*_ = −5*mV* for inhibitory synapses. Each presynaptic spike train was taken from a Poisson process with a constant instantaneous firing rate. Values used *f*_*exc*_ = 3.3*Hz* for excitatory synapses and *f*_*inh*_ = 3.2*Hz* for inhibitory synapses. Resulting output average firing rate for these simulation values was 2.1Hz.

### L5PC simulation

For Fig. 2 and Fig. 3 simulations, we used a detailed compartmental biophysical model of cortical L5PC **as is**, modeled by Hay et. al, 2011. For full description of the model please see **Methods** in the original paper. Briefly, this model contains in total 12 ion channels for each dendritic compartment. Some of the channels are unevenly distributed over the dendritic arbor. In Fig. 3 double exponential conductances based AMPA synapses were used in simulations with *τ*_*rise*_ = 0.3*ms*, *τ*_*depress*_ = 3*ms* and *g*_*max*_ = 0.4*ns*. For Fig. 2 and Fig. 4, in related simulations we used the standard NMDA model by Jahr and Stevens, 1993, with *τ*_*rise*_ = 2*ms*, *τ*_*depress*_ = 70*ms*, *γ* = 0.08 *mV*^−1^ and *g*_*max*_ = 0.4*ns*. For both Fig. 2 and Fig. 3, we also used double exponential GABA A synapses with *τ*_*rise*_ = 2*ms*, *τ*_*depress*_ = 8*ms* and *g*_*max*_ = 1*nS* on each independent dendritic segment, we placed a single AMPA (for Fig. 3) or AMPA+NMDA (for Fig. 2) synapse as well as a single GABA A synapse. In order to mimic uniform coverage of excitatory and inhibitory synapses on the entire dendritic tree, we stimulated each compartment with a firing rate proportional to the segment’s length. Each presynaptic spike train was taken from a Poisson process with a smoothed piecewise constant instantaneous firing rate. The Gaussian smoothing sigma, as well as the time window of constant rate before smoothing were independently resampled for each 6-second simulation from the range 10ms to 1000ms (Fig. S1**D**). This was chosen, as opposed to a constant firing rate, to create additional temporal variety in the data in order to increase the applicability of the results to all possible situations. For Fig. 2 simulations with NMDA synapses, the total amount of excitatory and inhibitory presynaptic spikes per 100ms range between 0 and 800 spikes (Fig. S1). This is equivalent to 8000 excitatory synapses with an average rate of 1Hz and 2000 inhibitory synapses with an average rate of 4Hz. The average output firing rate of the simulated cell across all simulations was 1.0Hz. For Fig. 3 simulations, with AMPA only synapses, the total amount of excitatory and inhibitory presynaptic spikes per 100ms range were increased in order to account for the smaller amount of total current injected due to lack of NMDA current, with the purpose to achieve similar output firing rates of 1.0Hz. See Fig. S8 for detailed comparison.

### L23PC simulation

For Fig. 4 simulations, we used a detailed compartmental biophysical model of cortical L23PC **as is**, modeled by Branco et. al, 2010. In these experiments we stimulated a single branch with 9 dendritic segments with an NMDA synapse on each compartment, with parameters as in the simulation for Fig. 4. The branch was selected as in Branco et al, 2010 in order to perform comparison with original paper. Similarly, to Fig. 2. and Fig. 3 simulations, each presynaptic spike train was taken from a Poisson process with a smoothed piecewise constant instantaneous firing rate. The number of presynaptic input spikes to the branch per 100ms ranged between 0 and 15 in simulations used for training. In Fig. 4**C**,4**E**,4**F**, we repeated input stimulation protocol suggested by Branco et. al, 2010, consisting of single presynaptic spike per synapse with constant time intervals of 5ms between subsequent synaptic activations, only randomly permuting the order of activation between trials.

### DNN fitting

In order to represent the input in a suitable manner for fitting with a DNN, we discretize time using 1ms time bin Δ*t*. Using this discretization, we can represent a spike train as a sequence of binary values *S*[*t*], such that *S*[*t*] ∈ {0,1}, since the length of a spike is approximately 1ms there cannot be more than a single spike in such a time interval. We denote the spike trains the neuron receives as input as *X*[*s*, *t*], *s* ∈ {1,2, …, *N*_*syn*_}, *t* ∈ {1,2, …, *T*}, where *s* denotes the synapse index, and *t* denotes time. The spike trains a neuron emits as output we denote as *y*_*spike*_[*t*], The somatic voltage trance we denote as *y*_*voltage*_[*t*]. For every point in time, we attempt to predict both somatic spiking *y*_*spike*_[*t*] and somatic voltage *y*_*voltage*_[*t*] based only a *T*_*input*_ sized window of presynaptic input spikes. i.e. define the vector 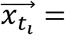 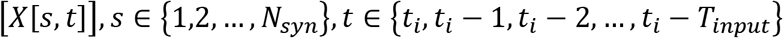 and a neural network that maps 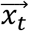 to 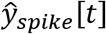 and 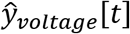. i.e. 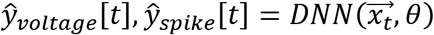. We treat spike prediction as a binary classification task and use standard log loss and treat voltage prediction as regression task and use standard MSE loss. We wish to find a model’s parameters *θ* such that we minimize a combined loss 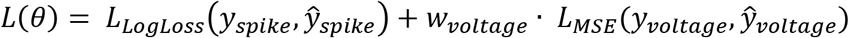, where *w*_*voltage*_ is the relative importance of the spike prediction loss with respect to the somatic voltage prediction loss. For most of our experiments we set *w*_*voltage*_ to be about half the size of the spike loss. The DNN architecture we used was a temporally convolutional network (TCN) (Bai, Zico Kolter, and Koltun 2019) and we applied it in a fully convolutional manner on all possible timepoints. Note that when the temporal filter size after the first layer is 1 in a TCN applied as described, this is effectively a fully connected neural network. In most of our experiments we used fully connected neural networks, except for Fig. 2 in which we used a proper TCN with hierarchical convolutional structure. After every convolutional layer, a batch normalization layer immediately follows. We employed a learning schedule regime in which we lowered the learning rate and increased batch size as we progressed through training. Full details of the learning schedule in each case are in the attached code repository. For generation of Fig. 5S. we trained many networks with different hyperparameters and trained each network for 2-7 days on a GPU cluster consisting of several V100, K80 and 2080Ti Nvidia GPUs. All results of the different hyperparameters and results can be found in the data link on the Kaggle platform.

### Model evaluation

We divided our simulations to train, validation and test datasets. We fitted all DNN models on train dataset, and all reported results are on an unseen test dataset. A validation dataset was used for modeling decisions, hyperparameter tuning and snapshot selection during the training process (early stopping). We evaluated binary spike prediction results using receiver operator characteristic (ROC) curve and calculated the area under curve (AUC). We note that due to the relatively low firing rate of the neuron, the binary classification problem of instantaneous spike prediction problem is highly unbalanced. For every second of simulation there was on average 1 positive sample (spike) for every 999 negative samples (non-spikes). Therefore, we used a very conservative threshold over the binary spike probability prediction output of the DNN in order to create the final spike train prediction and examine the cross-correlation plot. Note also that a prediction without a single True Positive on the 1ms time binning binary spike prediction problem can still be in fact a very good solution, e.g., if our model outputs, as its prediction, the exact same spike train as the original but offset by 1ms in time. In this case, there will be no True positives, and many False positives, but the predicted spike train is quite good nonetheless, namely: the temporal cross correlation between the original and predicted spike trains is not directly related to binary prediction metrics used and therefore we display it. For creating a binary prediction, we chose a threshold that corresponds to 0.2% false positive rate. In order to evaluate the temporal precision of the binary spike prediction we plotted the cross-correlation between the predicted output spike train and the ground truth simulated spike train. In order to evaluate the voltage prediction, we calculated the RMSE and plot the scatter plot between predicted voltage and the ground truth simulated voltage.

## Code and data availability

Data and pretrained networks that were used in this work are available on Kaggle at the following link: https://www.kaggle.com/selfishgene/single-neurons-as-deep-nets-nmda-test-data.

A short python script that loads a pretrained artificial network and makes a prediction on the entire NMDA test set that replicates the main result of the paper (Fig. 2) can be found in the following link: https://www.kaggle.com/selfishgene/single-neuron-as-deep-net-replicating-key-result.

A short python script that loads the data and explores the dataset (Fig. S1 & Fig. S2) can be found in the following link: https://www.kaggle.com/selfishgene/exploring-a-single-cortical-neuron.

## Acknowledgements

We thank Oren Amsalem, Guy Eyal, Michael Doron, Toviah Moldwin, Yair Deitcher, Eyal Gal and all lab members of the Segev and London Labs for many fruitful discussions and valuable feedback regarding this work. This work was supported by a grant from the EU Horizon 2020 program (720270, Human Brain Project), a grant from the Gatsby Charitable Foundation and a grant by Huawei Technologies Co. Ltd.

## Supplementary Figures

**Fig S1.**
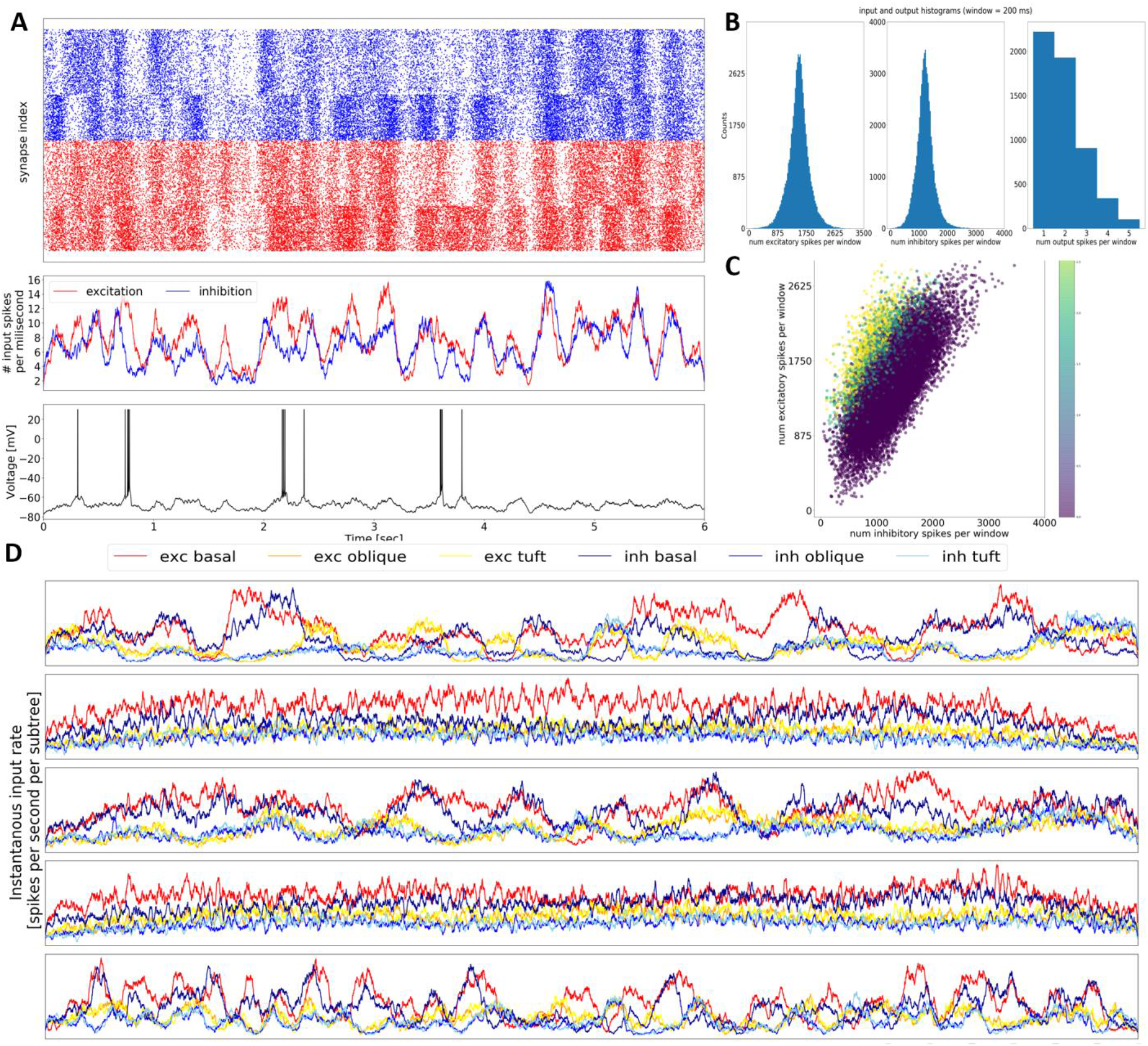
Simulations Input-Output of an *in vivo* like regime of a detailed L5PC neuron model with NMDA synapses. (**A**) Inputs and outputs of a sample simulation. Top, raster plot of presynaptic input excitatory spikes (red) and inhibitory spikes (blue). Middle, total number of excitatory spikes per 1 millisecond. Smoothed with a Gaussian kernel with tau=20ms. Bottom, somatic voltage trace and spiking output of the same simulation (**B**) Histograms of number of excitatory input spikes (left), inhibitory input spikes (middle) received as input on the entire dendritic tree in a 200ms time window for the L5PC model with NMDA synapses. Right. Histogram of number of output spikes in the same time period. (**C**) Scatter plot expanding the information presented in (B): horizontal axis is number of inhibitory spikes, vertical axis is the number of excitatory spikes and the color of each dot represents the number of output spikes for the same time window of 200ms. (**D**) Example illustration of inputs, in units of spikes per millisecond per subtree and synapse type, for 5 different simulations. We can see a wide amount of variability in input regimes of our simulations.

**Fig S2.**
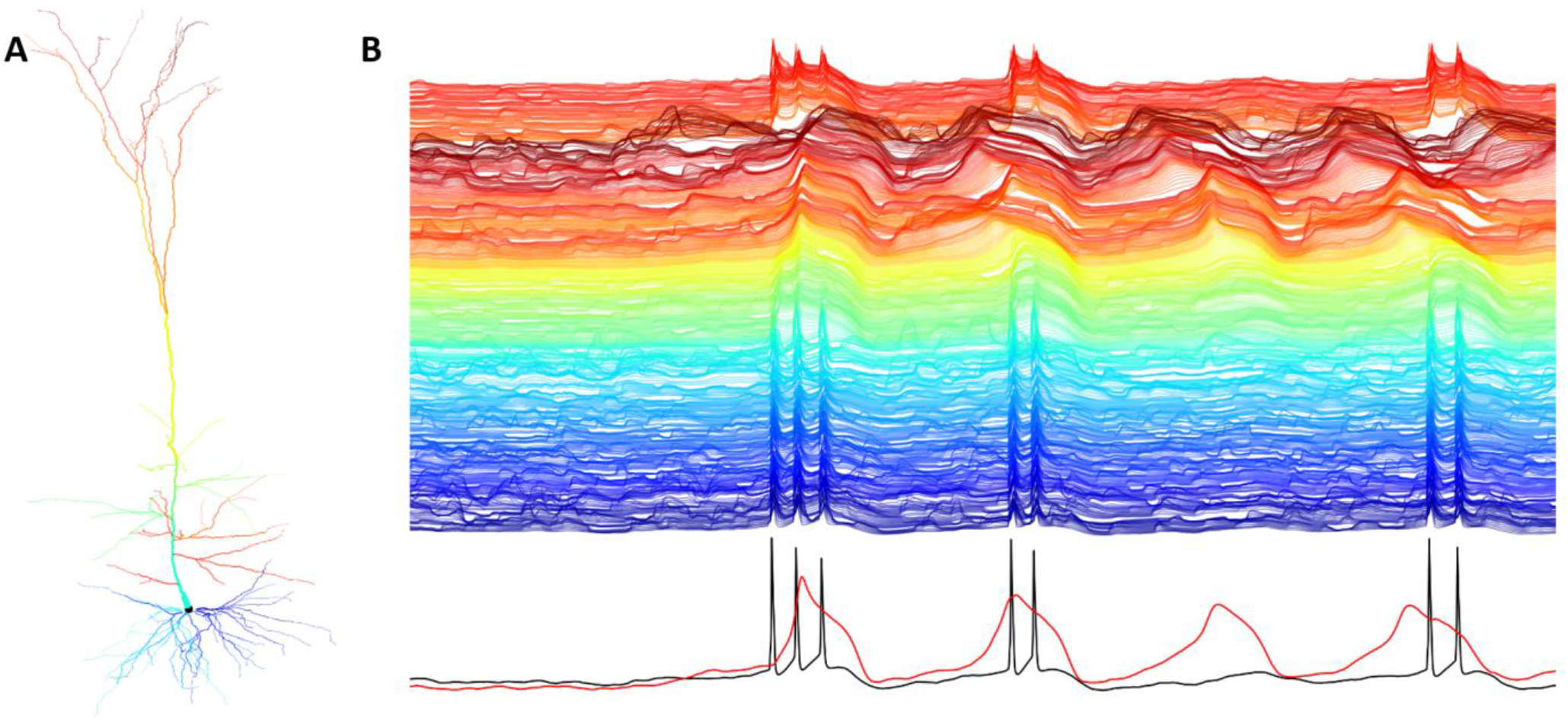
Dendritic voltage traces of *in vivo* like input regime of a detailed L5PC neuron model with NMDA synapses. (**A**) Illustration of the simulated cell morphology. (**B**) Top. Local dendritic voltage traces of all 639 simulated compartments. The colors of each trace are color-coded as the colors of the morphology illustration in (A). Bottom. Somatic (black) and nexus (red) voltage traces.

**Fig S3.**
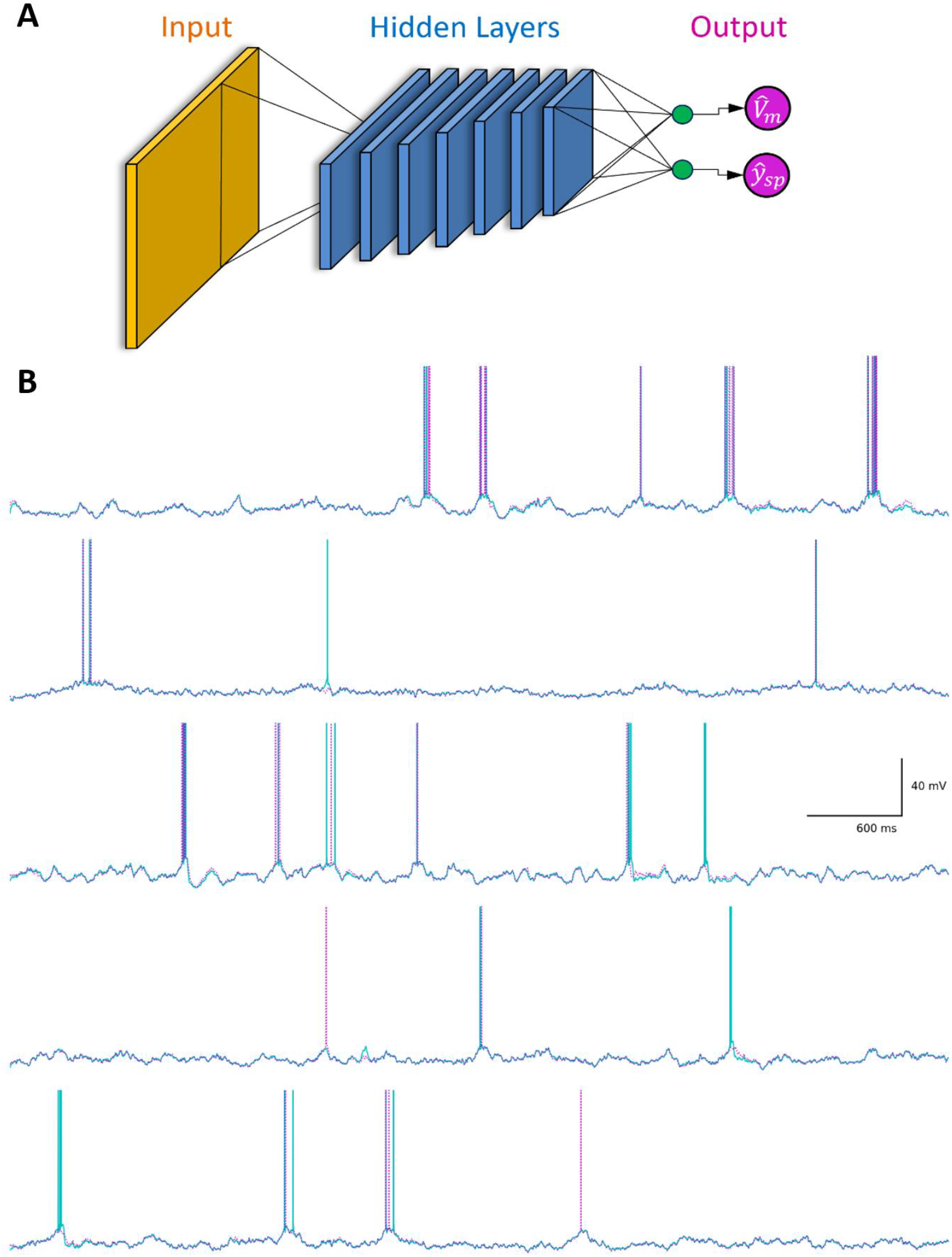
Additional ground truth vs prediction traces of a large TCN mimicking an L5PC neuron with NMDA synapses (Fig. 2). (**A**) Illustration of TCN architecture used to predict somatic voltage and spiking of L5PC detailed simulation. Same as Fig. 2 (**B**) Five randomly selected exemplar voltage responses of the L5PC model with NMDA synapses (cyan) and of the analogous DNN (magenta) to random synaptic input stimulation.

**Fig S4.**
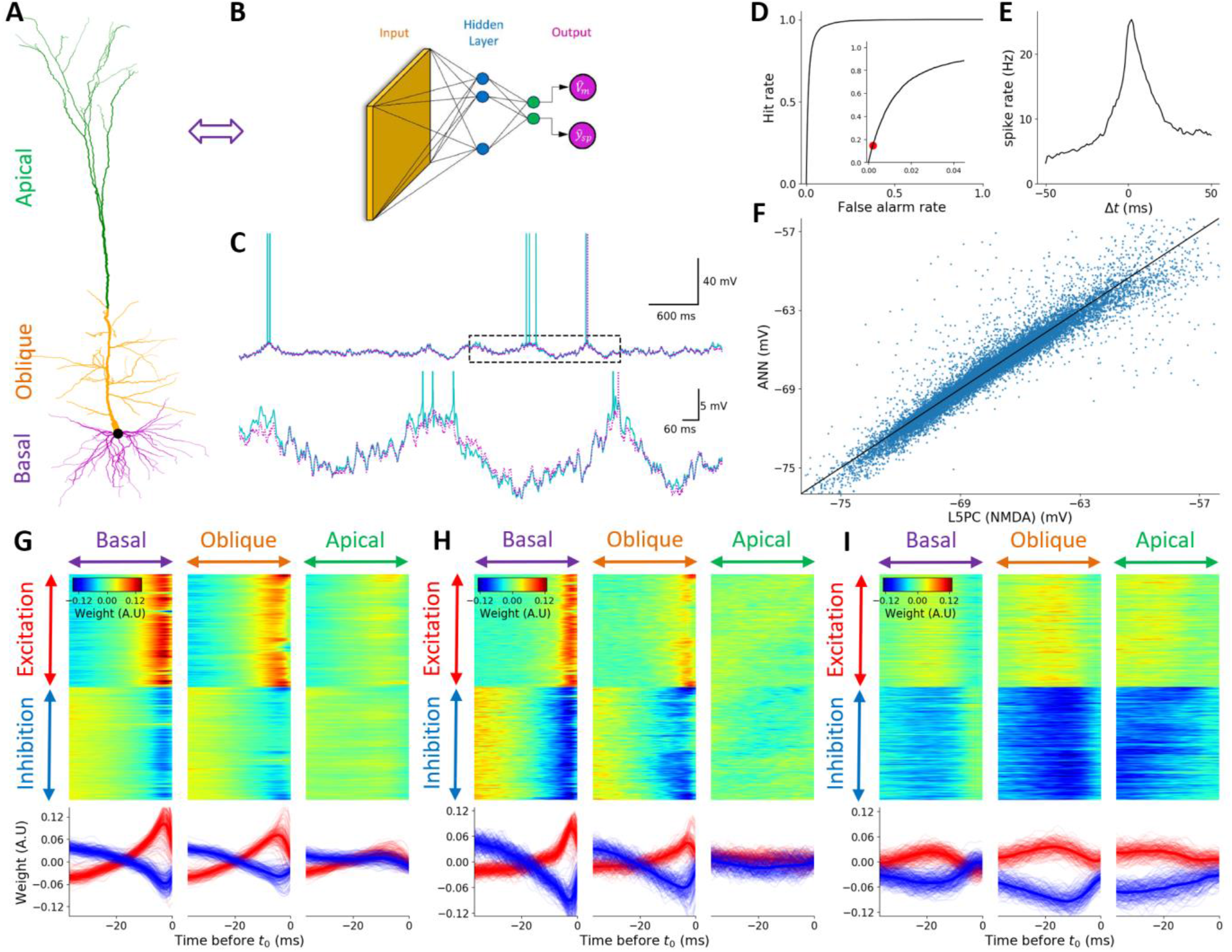
Detailed L5PC neuron model with NMDA synapses is not captured by a FCN with 1 hidden layer consisting of 128 hidden units at each layer. (**A**) Illustration of L5PC model. Basal oblique and apical dendrites are marked by respective purple, orange and green colors. (**B**) analogous DNN. Orange, blue and magenta circles represent the input layer, the hidden layer and the DNN output, respectively. Green units represent linear activation units. (Detailed architecture: 1 hidden layer, 128 units per layer, time window extent of 43ms) (**C**) Top. Exemplar voltage response of the L5PC model with NMDA synapses (cyan) and of the analogous DNN (magenta) to random synaptic input stimulation. Bottom. Zoom in on the dashed rectangle region in the top trace. (**D**) ROC curve of spike prediction; the area under the curve (AUC) is 0.9769. Zoom in on up to 4% false alarm rates is shown in the inset. Red circle denotes the threshold selected for the DNN model shown in **B**. (**E**) Scatter plot of the predicted DNN subthreshold voltage versus ground truth voltage. (**F**). Cross Correlation plot between the ground truth (L5PC with NMDA synapses) spike train and the predicted spike train of the respective DNN, when the prediction threshold was set according to red circle in **D**. (**G**) Learned weights of a selected unit in the DNN. Top Left, top center and top right, inputs located on the basal dendrites, on the oblique dendrites, and on the apical tuft, respectively. For each case, top half of the rows are excitatory synapses whereas bottom half of rows are its inhibitory synapses. Different columns correspond to different time points relative to t_0_ (rightmost time point). Bottom. temporal cross-section of the learned weights above. (**H**) Similar to **G**, first layer weights for a different unit in the first layer but with a different spatio-temporal pattern. (**I**) An additional unit, similar to **G** & **H**.

**Fig S5.**
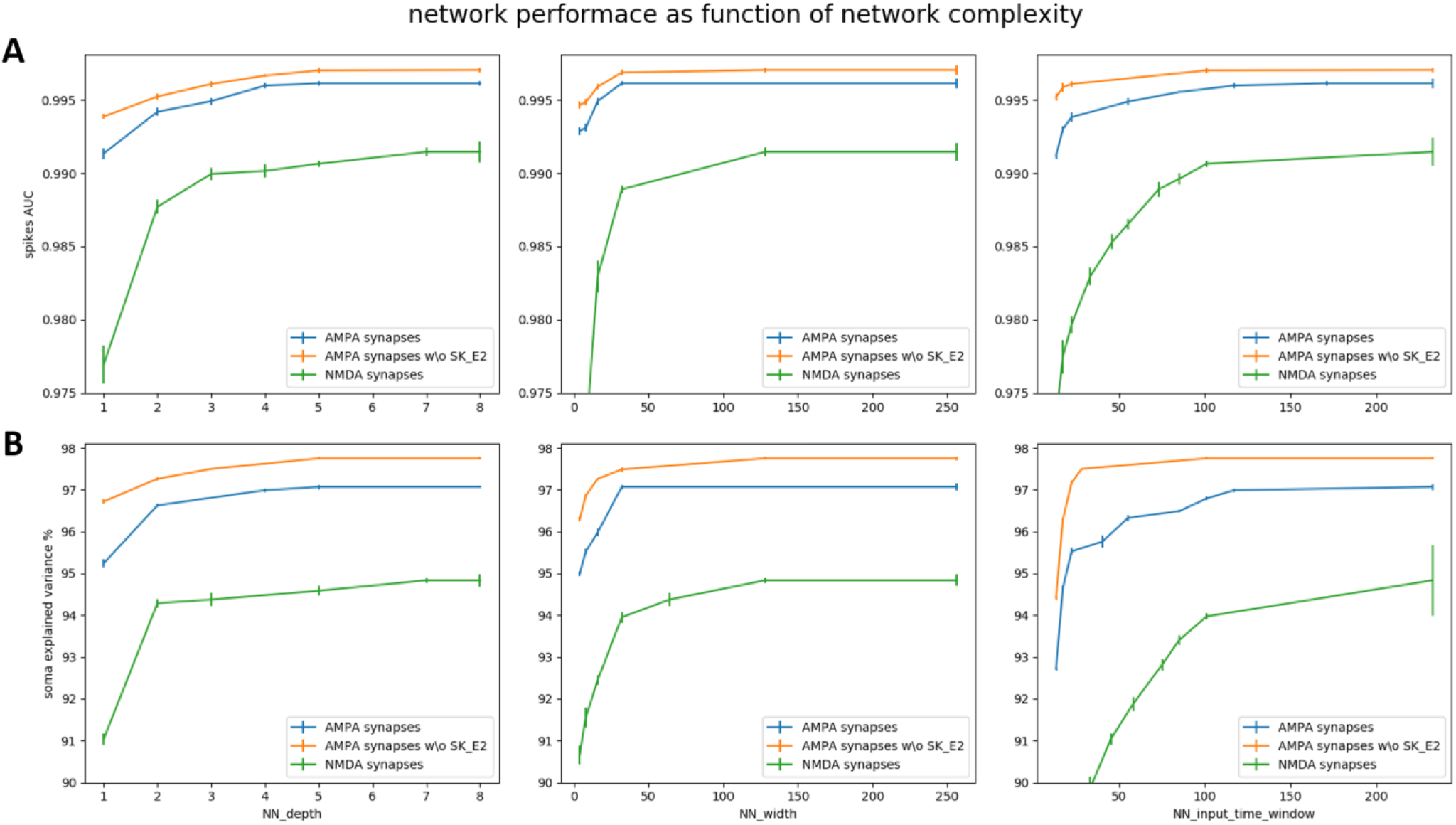
The complexity of DNNs required to fit L5PC with NMDA synapses is substantially greater than L5PC with AMPA only synapses. (**A**) Left, spiking prediction accuracy measure (AUC) as function of the depth of the fitted DNN for 3 different cases. Green curve is full biophysical model with NMDA synapses, blue curve is full biophysical model with AMPA only synapses, and orange curve is similar to the AMPA only case but with an additional SK ion channel removed from the soma. Variance bars are across different test data subsets. Middle, similar plot to the one on the left but for the width of the DNN (i.e., number of channels/feature maps). Right, like previous two plots, but for the temporal extent of the input time window (i.e., effective “memory” of the neuron) (**B**) Similar plots in **A**, but now the vertical axis represents the somatic subthreshold prediction accuracy (as depicted by percent of variance explained). <u>Note</u>: each point of each of the individual curves represents the **best** network across all other variables in that figure.

**Fig S6.**
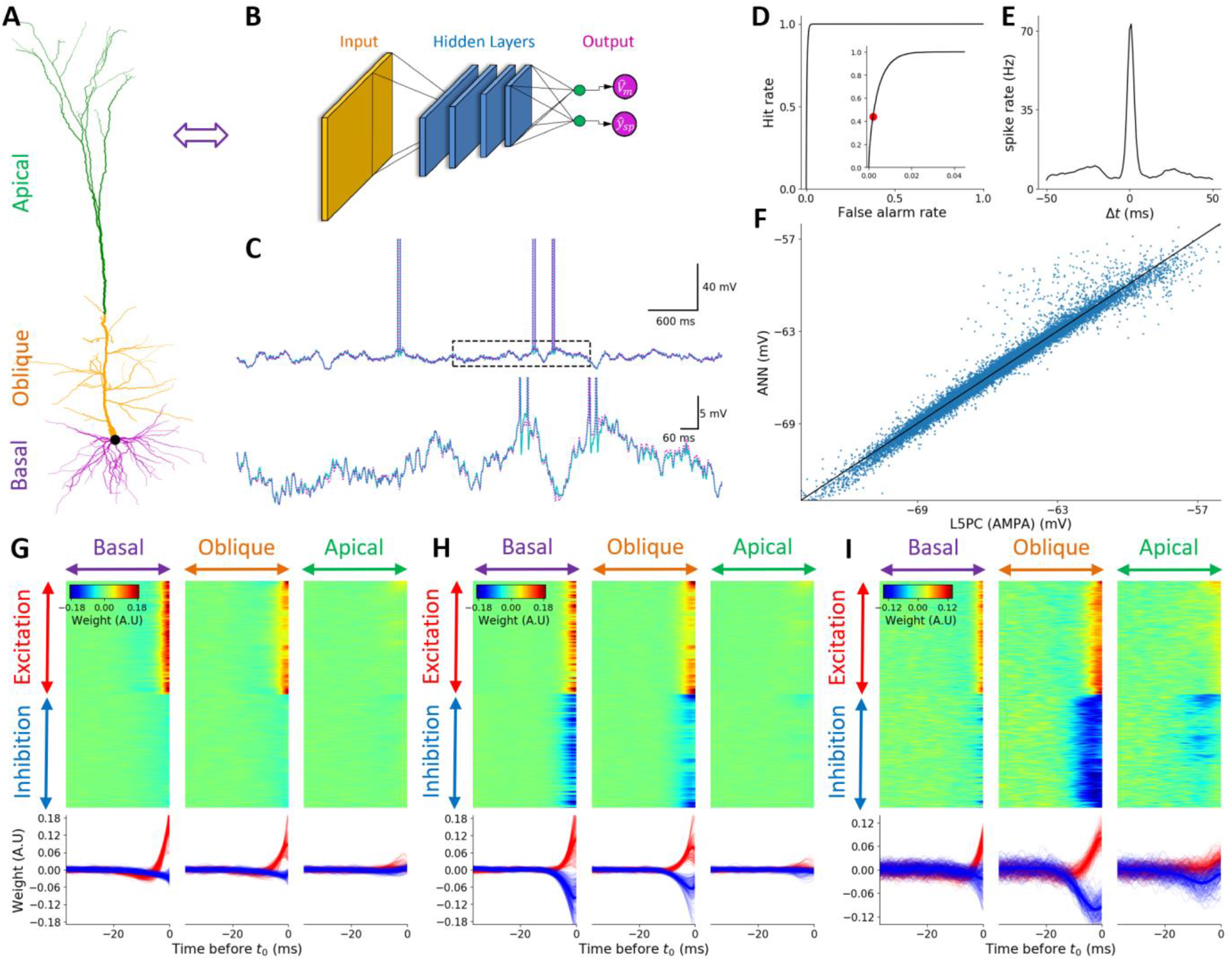
Moderately large TCN fits detailed L5PC neuron model with AMPA synapses with an extremely high degree of prediction accuracy. (**A**) Illustration of L5PC model. Basal, oblique, and apical dendrites are marked by purple, orange and green colors, respectively. (**B**) Analogous DNN. Orange, blue, and magenta circles represent the input layer, the hidden layer, and the DNN output, respectively. Green units represent linear activation units. (detailed architecture: 4 temporally convolutional layers, 64 features maps per layer, time window extent of 120ms) (**C**) Top. Exemplar voltage response of the L5PC model with AMPA synapses (cyan) and of the analogous DNN (magenta) to random synaptic input stimulation. Bottom. Zoom in on the dashed rectangle region in the top trace. (**D**) ROC curve of spike prediction; the area under the curve (AUC) is 0.9959. Zoom in on up to 4% false alarm rates is shown in the inset. Red circle denotes the threshold selected for the DNN model shown in **B**. (**E**) Scatter plot of the predicted DNN subthreshold voltage versus ground truth voltage. (**F**). Cross Correlation plot between the ground truth (L5PC with AMPA synapses) spike train and the predicted spike train of the respective DNN, when the prediction threshold was set according to red circle in **D**. (**G**) Learned weights of a selected unit in the DNN. Top Left, top center, and top right, inputs located on the basal dendrites, on the oblique dendrites, and on the apical tuft, respectively. For each case, top half of the rows are excitatory synapses whereas bottom half of rows are its inhibitory synapses. Different columns correspond to different time points relative to t_0_ (right most time point). Bottom. temporal cross-section of the learned weights above. (**H**) Similar to **G**, first layer weights for a different unit in the first layer but with a different spatio-temporal pattern. (**I**) An additional unit, similar to **G** & **H**.

**Fig S7.**
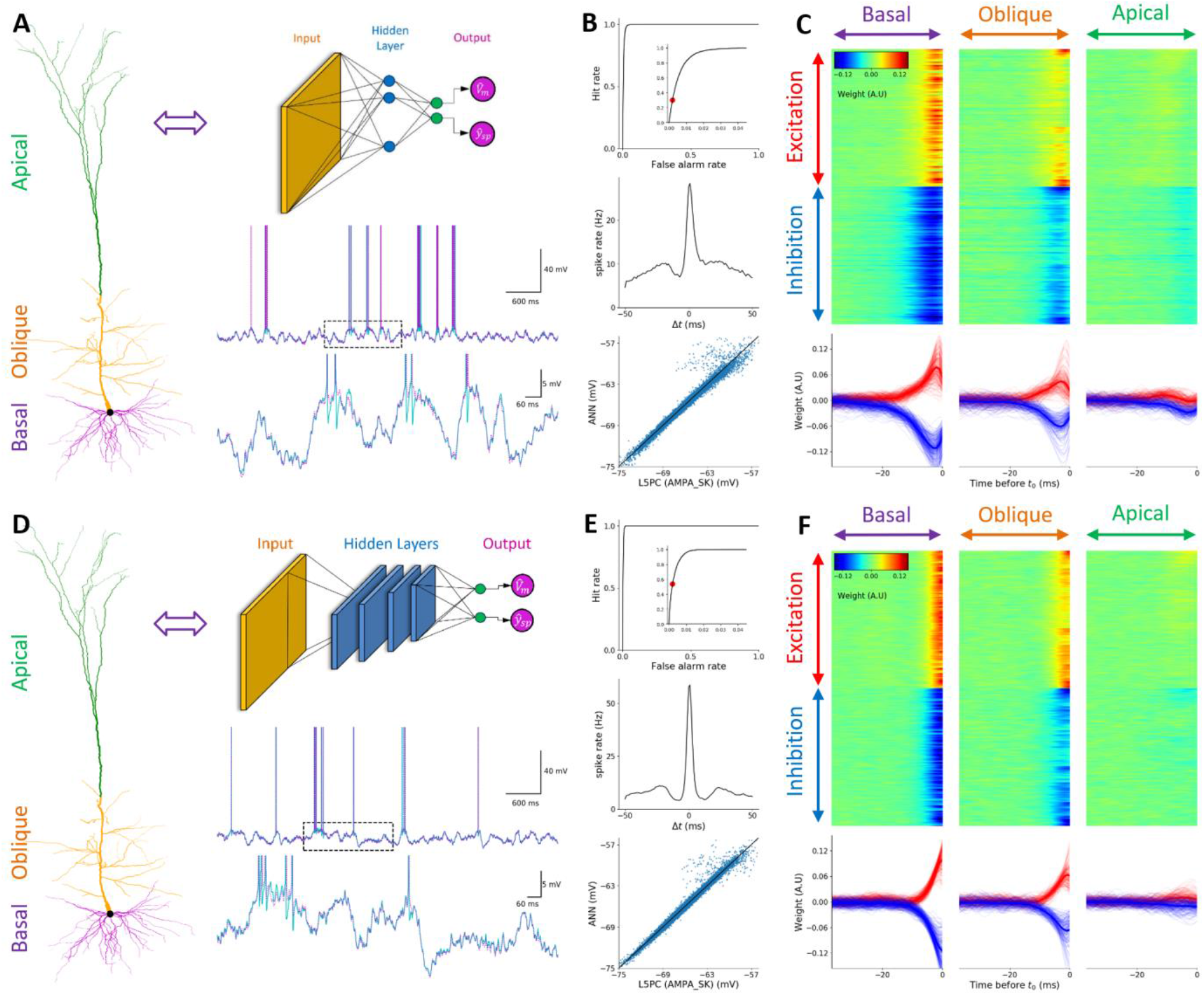
Removal of SK channels results in even higher prediction accuracy of both large and small DNNs. (**A**) DNN illustration of 1-layer FCN and exemplar traces of prediction. Like Fig. 3**A**,3**B**,3**C** (**B**) Quantitative performance evaluation of the network depicted in **A**, similar to Fig. 3**D**,3**E**,3**F**, only this time without SK channels in the cell. (**C**) Learned weights of a selected first layer unit of the network depicted in **A**. (**D**) DNN illustration of 4-layer TCN and exemplar traces of prediction. Similar to **B**(**E**) Quantitative performance evaluation of the network depicted in **D**. (**C**) Learned weights of a selected first layer unit of the network depicted in **D**.

**Fig S8.**
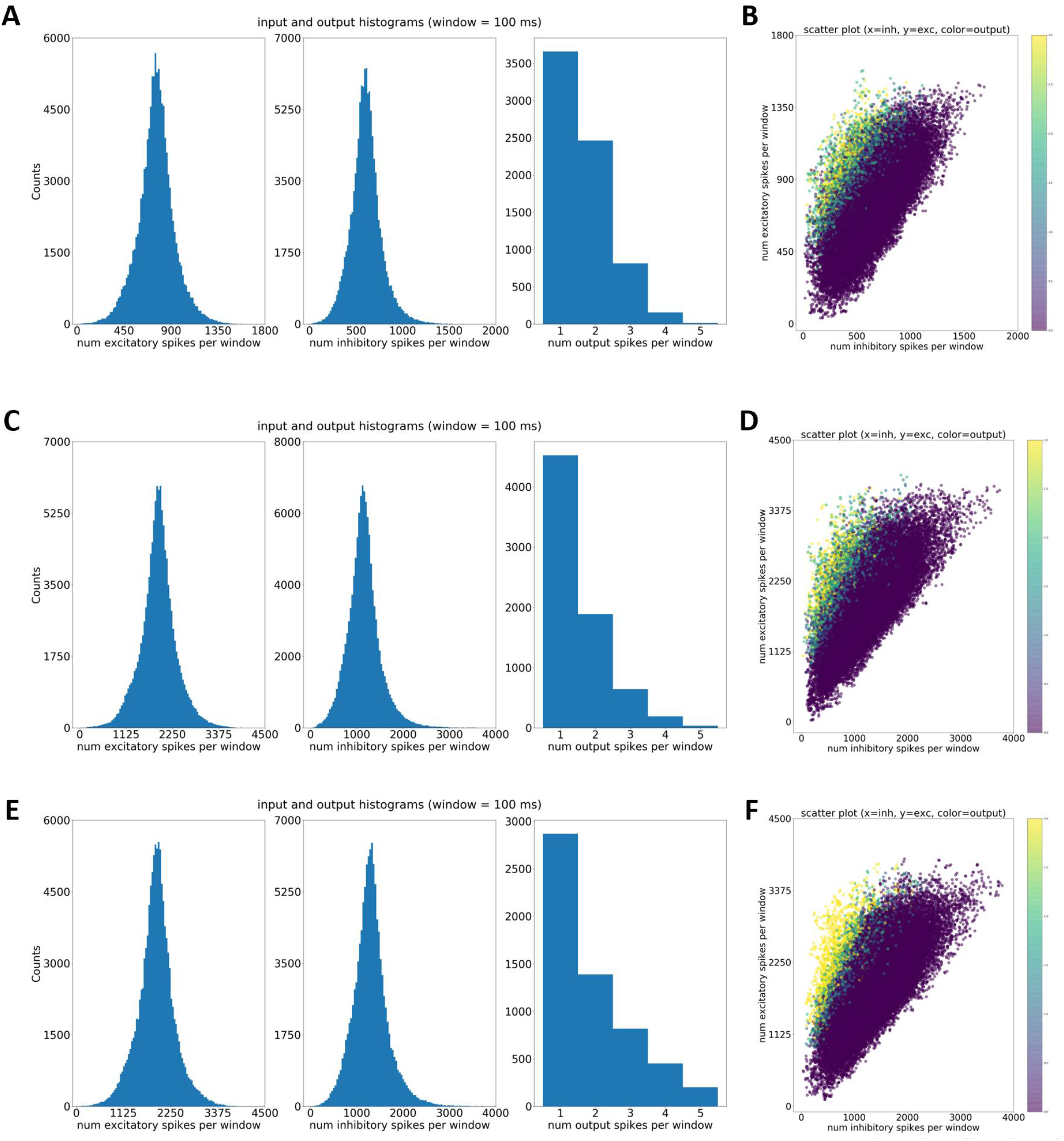
Input-Output regimes of detailed L5PC model in the 3 different simulation cases: NMDA synapses, AMPA synapses and AMPA synapses with an additional SK channel removed. (**A**) Histograms of number of excitatory input spikes (left), inhibitory input spikes (middle) received as input on the entire dendritic tree in a 100ms time window for the L5PC model with NMDA synapses. Right. Histogram of number of output spikes in the same time period. (**C**) Scatter plot expanding the information presented in **A**: horizontal axis is number of inhibitory spikes, vertical axis is the number of excitatory spikes and the color of each dot represents the number of output spikes for the same time window of 200ms. (**C-D**) Similar to **A-B**, but for the case of L5PC model with AMPA only synapses. (**E-F**) Similar to **A-B**, but for the case of L5PC model with AMPA only synapses and SK channels removed from the cell.

**Fig S9.**
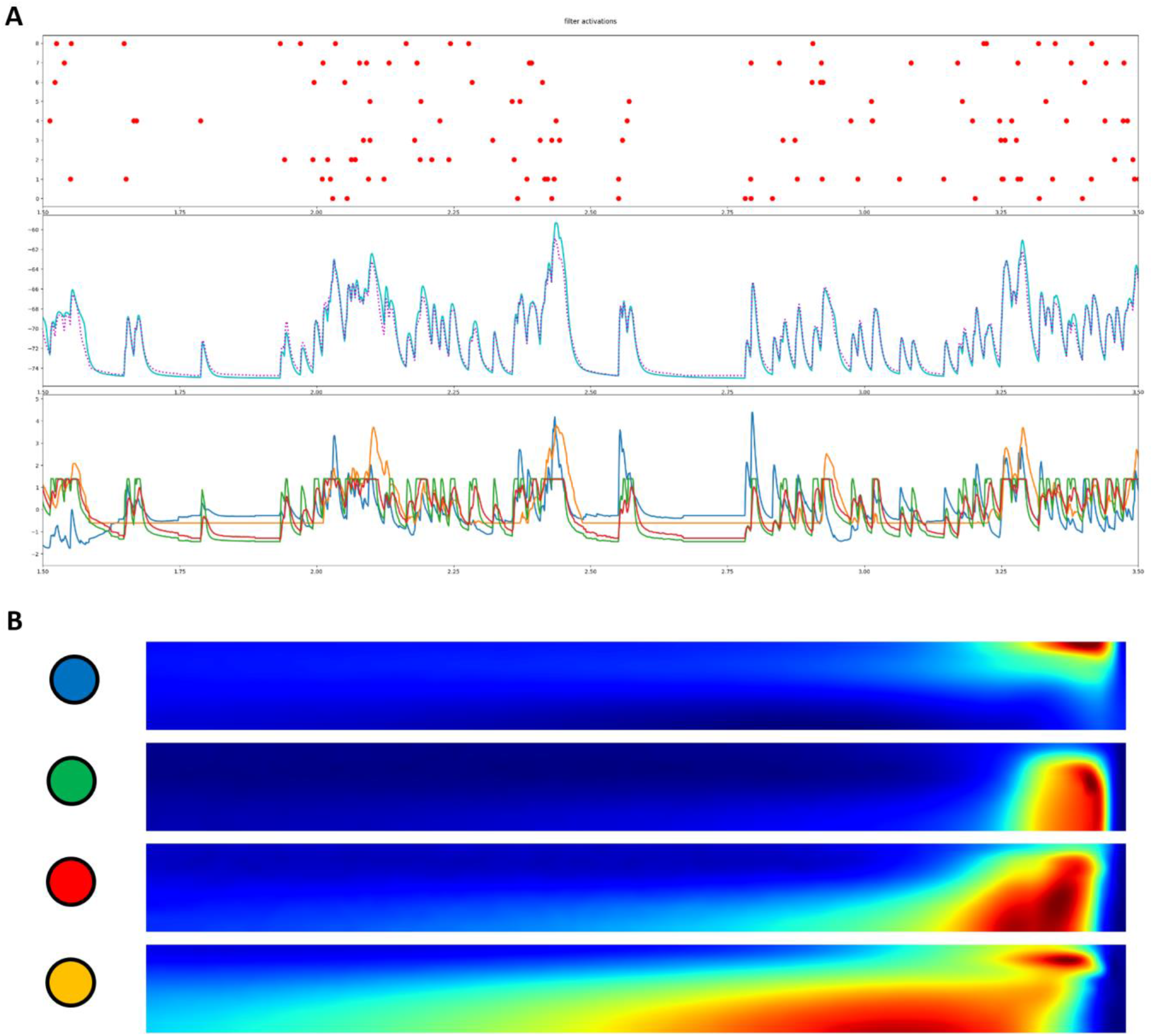
Interpretation of first layer filter activations of DNN representing a single dendritic branch. (**A**) Top. Raster plot of presynaptic input spikes to the branch from Fig. 4. Middle. Somatic voltage trace (cyan) and predicted voltage by the DNN from Fig. 4. (magenta). Bottom. Activations of the DNNs 4 units (**B**) Illustration of the DNN first layer weights and the colors they are represented by in the bottom plot of **A** (circles on the left of each filter).

## Notes

#### Summary of Updates

The updated text contains a small narrative change, a more precise discussion section, several additional relevant citations were added, and 9 supplementary figures were also added. More importantly, links to code data and pretrained models were added. In the updated work we used larger and more diverse training and testing datasets, we trained the models longer that resulted in better performing models, and performed more rigorous hyper parameter sweep.

